# Novel Approach Explains Spatio-Spectral Interactions in Raw Electroencephalogram Deep Learning Classifiers

**DOI:** 10.1101/2023.02.26.530118

**Authors:** Charles A. Ellis, Abhinav Sattiraju, Robyn L. Miller, Vince D. Calhoun

## Abstract

The application of deep learning classifiers to resting-state electroencephalography (rs-EEG) data has become increasingly common. However, relative to studies using traditional machine learning methods and extracted features, deep learning methods are less explainable. A growing number of studies have presented explainability approaches for rs-EEG deep learning classifiers. However, to our knowledge, no approaches give insight into spatio-spectral interactions (i.e., how spectral activity in one channel may interact with activity in other channels). In this study, we combine gradient and perturbation-based explainability approaches to give insight into spatio-spectral interactions in rs-EEG deep learning classifiers for the first time. We present the approach within the context of major depressive disorder (MDD) diagnosis identifying differences in frontal δ activity and reduced interactions between frontal electrodes and other electrodes. Our approach provides novel insights and represents a significant step forward for the field of explainable EEG classification.

## 1. INTRODUCTION

A growing number of studies have applied deep learning (DL) classifiers to raw resting-state electroencephalography (rs-EEG) data. Relative to studies that apply traditional machine learning (ML) methods to extracted EEG features [1], [2], DL classifiers do not presume which features will be useful for classification and perform automated feature engineering. Unfortunately, DL methods also have reduced explainability, and explainability approaches developed for DL classifiers in other domains [3] are often unhelpful for rs-EEG. As such, many studies have begun developing rs-EEG explainability methods [4]–[9]. However, existing approaches do not give insight into spatio-spectral feature interactions (i.e., they do not explain how frequency bands in one channel interact with activity in other channels), which are often used in rs-EEG analysis. In this study, we present, what is to our knowledge, the first explainability approach for raw rs-EEG classification that provides that insight by combining a gradient-based explainability approach with spatial and spectral perturbation. We demonstrate our approach within the context of automated major depressive disorder (MDD) diagnosis and find that individuals with MDD tend to have changes in frontal δ activity and have less interactions between frontal electrodes and other electrodes.

Traditional ML methods have been applied to electrophysiology data for many years. They typically use manual feature engineering. Common single-channel EEG features include spectral power [1], [2], [10], and common multichannel features include temporal and spectral connectivity [11], [12]. While ML methods can be effective, their use of manually engineered features inherently limits the feature space from which they can learn. In contrast, DL methods can perform automated feature extraction, which does not involve any a priori assumptions of what features will be useful. As such, as the field of DL has grown in prominence in recent years, more studies have begun applying DL approaches to rs-EEG data.

Unfortunately, rs-EEG DL methods also tend to be less explainable than ML approaches, and explainability approaches originally developed for DL models [3] in areas like image classification are often unhelpful for time-series classification. For example, a method like layer-wise relevance propagation (LRP) [3] may be directly translated to rs-EEG classification to indicate the importance of each time point to the classification of a given sample. However, when there are thousands of samples in a dataset and no way to align explanations temporally across samples, the resulting explanations may not be very helpful. Because of this, there has been a growth in studies presenting approaches for insight into spatial importance [4], spectral importance [5]–[7], and temporal waveform importance [5], [8], [9] in rs-EEG DL classifiers. Nevertheless, while great progress has been made in explaining rs-EEG DL classifiers, there are still gaps in existing capabilities. Namely, multi-channel features are often used to analyze EEG activity [11]–[13], but existing rs-EEG explainability approaches do not explain inter-channel interactions.

In this study, we present the first approach for raw rs-EEG DL classification that provides insight into the interaction between spectral activity in a specific channel with activity in other channels. Similar to existing approaches, it also identifies the importance of each channel and frequency band within each channel. It combines LRP [3] with a spectral perturbation approach. We implement the approach within the context of a one-dimensional convolutional neural network (1D-CNN) trained to differentiate between individuals with MDD (MDDs) and healthy controls (HCs). Importantly, many studies have examined MDD with rs-EEG [14]–[16], which creates a useful point of comparison for our explainability results. We find that MDDs have changes in frontal δ activity and have less interactions between frontal electrodes and other electrodes. These findings agree with previous studies and corroborate the viability of our proposed approach.

## 2. METHODS

In this section, we describe and discuss our approach.

### 2.1. Data Collection

We used a scalp EEG dataset [14] that is publicly available and has been used in multiple studies [15], [16]. It has 28 healthy controls (HCs) and 30 individuals with MDD (MDDs). All participants gave informed consent prior to data collection at the Hospital Universiti Sains Malaysia (HUSM) in Kelantan, Malaysia. Recordings were performed at resting state with eyes closed for 5-10 minutes at a sampling frequency of 256 Hertz. The standard 10-20 format with 64 electrodes was used for during recording.

### 2.2. Data Preprocessing

Due to the high levels of correlation present between scalp EEG channels, we reduced the size of the dataset to 19 channels: Fp1, Fp2, F7, F3, Fz, F4, F8, T3, C3, Cz, C4, T4, T5, P3, Pz, P4, T6, O1, and O2. We downsampled the data

### 2.3. Model Development

We adapted an architecture originally developed in [17] for schizophrenia classification. Figure 1 shows our model architecture. Relative to the original architecture in [17], we added a number of batch normalization layers and converted ReLU activation functions to ELU activation functions. Each sample input had dimensions of 5,000 time points x 19 features. We used a 10-fold stratified group shuffle split cross-validation approach to ensure that epochs from the same participant were not distributed across training, validation, and test groups within the same fold and that resulting performance estimates were reliable. Approximately, 80%, 10%, and 10% of samples were assigned to training, validation, and test sets in each fold, respectively. We used a data augmentation approach to double the size of the training sets. We produced a copy of the training data in each fold, and we added Gaussian noise with mean of zero and standard deviation of 0.7 to the copy. This approach has been used in a previous study [18]. We also used a class-weighted loss function to account for class imbalances in the training set. We used the Adam optimizer with a learning rate of 0.0075. We trained with a batch size of 128 samples for 35 epochs, using early stopping if the model validation accuracy did not improve after 10 consecutive epochs. We selected the model from the epoch with the highest balanced validation accuracy for testing. When evaluating test performance, we calculated the mean and standard deviation (SD) of the balanced accuracy (BACC), sensitivity (SENS), and specificity (SPEC) across folds. All convolutional and dense layers, except for the final dense layer that had Glorot normal initialization, were initialized with He normal initialization. The test data from the model with the highest overall test BACC was used for explainability analyses.

**Figure 1.**
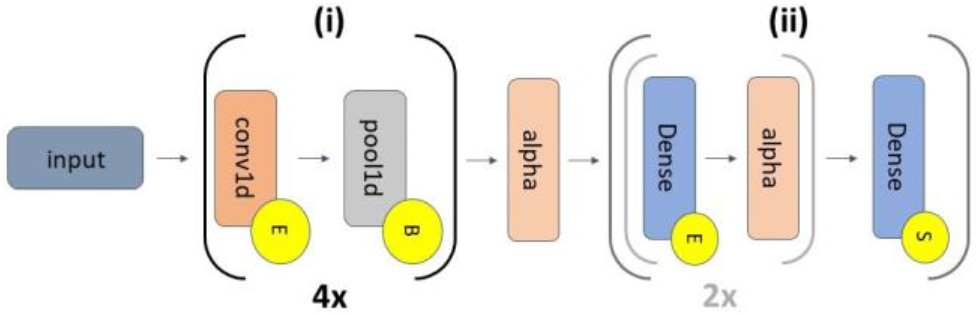
1D-CNN Architecture. The model has 2 sections that are separated by an alpha dropout layer (alpha): (i) feature extraction (which repeats 4 times) and (ii) classification. The grey inset within (ii) repeats twice. The 4 pairs of convolutional (conv1d) layer pairs have 5, 10, 10, and 15 filters, respectively (kernel size = 10, 10, 10, 5). The pairs are followed by a max pooling layer (pool size = 2, stride = 2). Component (ii) has 3 dense layers (64, 32, and 2 nodes). The first two dense layers have alpha dropout (alpha) with rates of 0.5. Additionally, the alpha dropout layer between (i) and (ii) has a rate of 0.5. ELU activations, batch normalization, and softmax activations are represented by yellow circles containing an “E”, “B”, or “S”. All conv1d and dense layers had max norm kernel constraints with a max value to 200 Hz. To increase the number of epochs available for training, we used a sliding window approach with a 2.5s step size to separate the data into 25-second epochs. We then channel-wise z-scored each participant separately. Our final dataset had 2,950 SZ and 2,942 HC epochs.

### 2.4. Explainability Approach

We combined LRP with a perturbation approach for insight into (1) the importance of each channel, (2) the importance of frequency bands in each channel, and (3) the interaction of frequency bands in each channel with other channels.

LRP is a popular explainability approach that has been applied in many neuroinformatic time-series analyses. It involves (1) forward passing a sample through a model, (2) assigning a total relevance value of 1 to the output node corresponding to the class of interest, and (3) backpropagating the total relevance back through the model to the input space with a relevance rule such that the total relevance at each layer still sums to 1. Relevance can be positive (i.e., providing evidence for a sample belonging to the class of interest) or negative (i.e., providing evidence for a sample belonging to classes other than the class of interest). We applied the αβ-rule (α=1, β=0) [3] to only propagate positive relevance.

Our composite explainability approach involved several steps. (1) We output relevance for each sample corresponding to its true class. After outputting relevance for each sample, we normalized the absolute relevance to force it to sum to 1 and summed the relevance associated with each of *C* channels. (2) We perturbed a frequency band *f* within a channel *c* by converting samples to the frequency domain with a Fast Fourier Transform (FFT), zeroing out Fourier coefficients corresponding to the frequency band *f*, and converting the perturbed sample back to the time domain with an inverse FFT. (3) We output the normalized and summed relevance associated with each of *C* channels for the perturbed samples. (4) We calculated the change in relevance assigned to each of *C* channels following the perturbation of frequency band *f* in channel *c*. (5) We repeated steps 2 through 4 for each of *F* frequency bands in *C* channels. Canonical frequency bands used in the analysis were δ (0 – 4 Hz), θ (4-8 Hz), α (8 – 12 Hz), β (12 – 25 Hz), γ_low_ (25 – 55 Hz), and γ_high_ (65 – 125 Hz).

Step 1 provided a spatial estimate of importance (i.e., the importance of each channel) to the classification. By analyzing the change in relevance in channel *c* following the perturbation of *F* frequency bands in channel *c*, we were able to gain insight into the relative importance of those *F* frequency bands in each channel (i.e., spatial + spectral importance). By analyzing the change in relevance in channels other than channel c following the perturbation of *F* frequency bands in channel *c*, we were able to gain insight into the interactions captured by the model between frequency bands in one channel and other channels. If the relevance of a particular channel decreased following the perturbation of channel *c*, it would follow that there was information in that particular channel that was only useful in the presence of frequency band *f* in channel *c*. If the relevance of a particular channel increased, that gives insight into how the model compensated for loss of information from frequency band *f* in channel *c*.

## 3. RESULTS AND DISCUSSION

In this section, we describe and discuss our findings along with limitations and future directions for our approach.

### 3.1. Model Performance

Table 1 shows the mean and SD of our model performance across folds and the performance for the test fold to which we applied our explainability approach. Our model performance is well-above chance-level. Additionally, the model on average prioritizes sensitivity over specificity. In the fold to which we applied our explainability approach, model test performance is at 100% across metrics.

**Table 1.** Model Performance Results

|  | SENS | SPEC | BACC |
| --- | --- | --- | --- |
| **Mean** | 81.15 | 71.06 | 76.11 |
| **SD** | 21.38 | 31.51 | 19.54 |

### 3.2. Spatial Importance

The top row of Figure 2 shows the relative importance of each channel to the model performance for each class. Across both HC and MDD groups, F7 tends to be highly important. Additionally, C3, Cz, and C4 also tend to be highly important across classes, and O1 and O2 tend to be very unimportant. However, there also tend to be class-specific differences in importance across channels. Interestingly, the model tends to prioritize Fp1 and Fp2 for MDDs and P4 and T6 for HCs.

**Figure 2.**
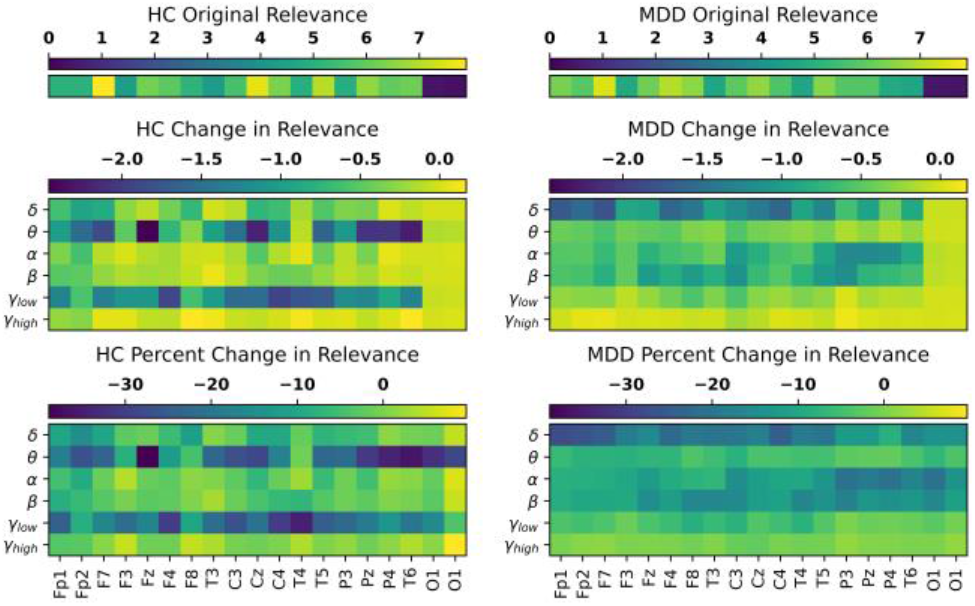
Spatio-spectral Importance. The top, middle, and bottom rows show the relevance of each channel, the change in relevance after the perturbation of each frequency band in each channel, and the percent change in relevance after the perturbation of each frequency band in each channel. HCs and MDDs are shown in the left and right columns, respectively. The y-axes show channels, and the x-axes show channels. The color bar above each panel shows the values associated with each heatmap. Each row of panels has identical color bars.

### 3.3. Spatial and Spectral Importance

The bottom two rows of Figure 2 show the relative importance of each channel-frequency band pair to the classifier. Interestingly, key frequency bands tend to be highly important across the majority of channels, rather than only being important for a subset of channels. There are class-specific differences in importance. The model prioritizes δ across most electrodes for MDDs, but δ is most important for frontal and frontoparietal electrodes, which fits with previous studies [14] that identified reduced δ activity in MDDs. The model also prioritizes α and β for MDDs across channels. In contrast, the model prioritizes θ and γ_low_ across channels for HCs. Previous studies have also found widespread effects of MDD upon θ [15].

### 3.4. Spatial and Spectral Interaction

Figure 3 shows the results for our interaction explainability approach. Interactions identified by the model are highly class-specific. The model identified widespread Fp1 and Fp2 interactions across channels for HCs but not for MDD, which is interesting given the importance of Fp1 and Fp2 for identifying MDDs. Additionally, Fz, F4, and Pz have interactions with most other electrodes in HCs but not MDDs. These findings fit with previous studies that have uncovered effects of MDD upon frontal connectivity [15]. In contrast, the model seems to have uncovered O1 interactions with most other electrodes for MDDs, though the importance of these interactions is questionable given the relative unimportance of O1 and O2 electrodes. Central electrodes interactions with other channels tend to be present for both classes. However, Cz and C4 interactions are more present in HCs than MDDs, and C3 interactions are more present in MDDs than HCs.

**Figure 3.**
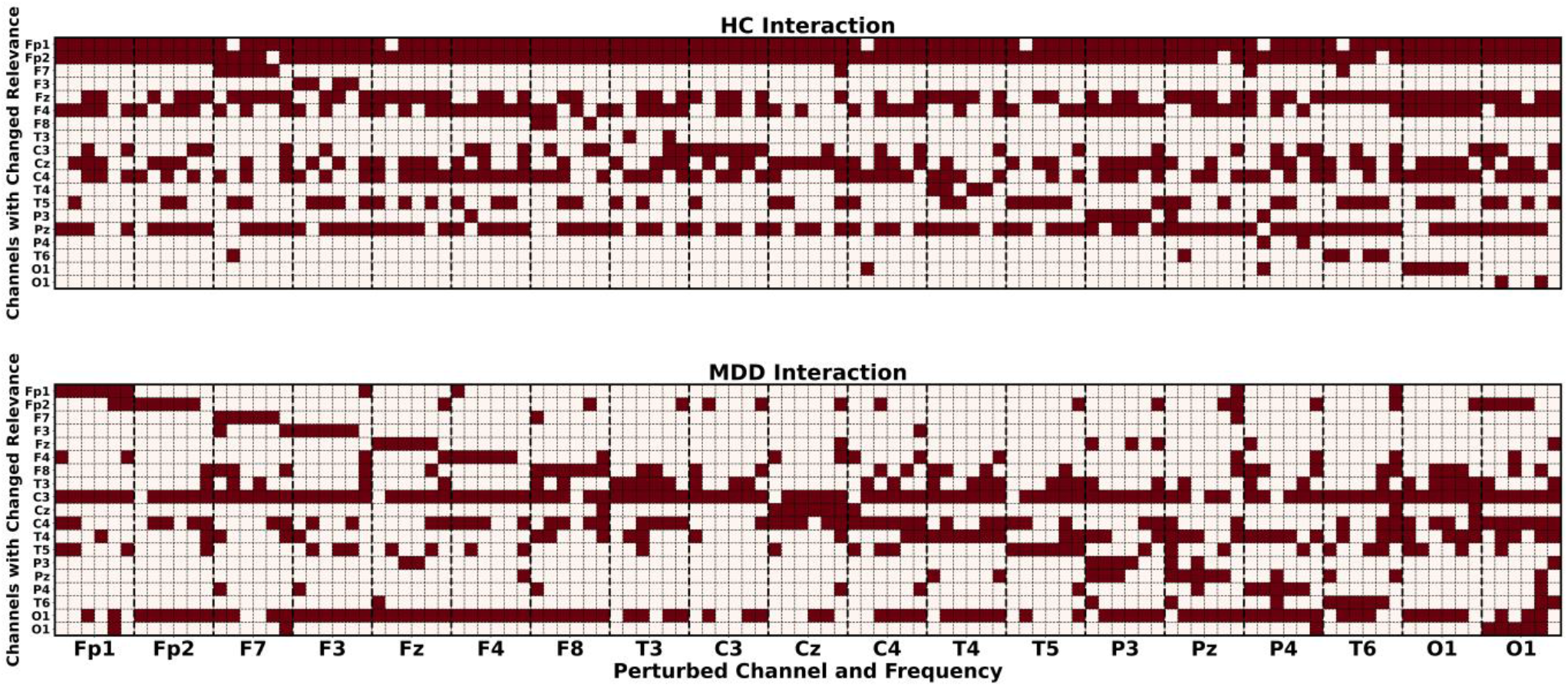
Spatio-Spectral Interactions. HC and MDD interactions are shown in the top and bottom panels, respectively. The x-axis shows perturbed channels and frequency bands. Vertical dashed lines separate values for each perturbed channel. There are 6 values associated with each perturbed channel that from left to right are δ, θ, α, β, γ_low_, and γ_high_. The y-axis indicates channels for which a change in relevance in response to the perturbed channel and frequency are being measured. Red values indicate the presence of a negative percent change in relevance.

### 3.5. Limitations and Next Steps

Our model performance is well-above chance-level and illustrates the viability of our explainability approach. However, our architecture and training approach could be further improved to enhance model performance. Additionally, while our approach offers a new capability for insight into spatio-spectral interactions, it can be improved. In future work, we plan to include statistical testing that identifies significant interactions. Our interaction approach is also likely not as robust as an approach like SHAP. However, estimating feature interaction in a sequential multi-feature perturbation or exhaustive manner like SHAP can be computationally prohibitive for the EEG space given the need to convert to and from the frequency domain. Additionally, an alternative feature interaction approach might examine the effect of perturbation on output layer activations. However, it is not uncommon for EEG models to have very high activations that do not greatly change in response to perturbation. Our use of the change in channel-specific relevance allows us to gain higher resolution insight into the effects of perturbation that can avoid this issue.

## 4. CONCLUSION

The use of deep learning models for raw rs-EEG classification is continuing to grow. However, this growth has also introduced problems related to explainability, which is particularly important for healthcare-related domains. Existing raw rs-EEG explainability approaches can give insight into the importance of different frequency bands or waveforms. However, an explainability approach capable of providing insight into the interaction of frequency bands in each channel with other channels has not yet been developed. This capability could be highly useful given the great importance of connectivity features to traditional EEG analyses. In this study, we present the first approach capable of giving insight into spatio-spectral interactions in multichannel rs-EEG data. We implement the approach within the context of MDD classification, identifying effects of MDD upon frontal δ activity and upon frontal interactions. Our approach represents a significant step forward for the domain of rs-EEG explainability and has potential applications across a variety of neurological and neuropsychiatric disorders. Moreover, we hope that it will encourage the field to continue the development of explainability approaches that give insight into multi-channel rs-EEG interactions.

## 5. ACKNOWLEDGMENTS

This research is supported by NIH R01MH123610, NIH R01MH118695, and NSF 2112455.

## REFERENCES

[1] N. Ince, F. Goksu, G. Pellizzer, A. Tewfik, and M. Stephane, “Selection of spectro-temporal patterns in multichannel MEG with support vector machines for schizophrenia classification.,” inProceedings of the 30th Annual International Conference of the IEEE Engineering in Medicine and Biology Society, 2008, pp. 3554–7.

[2] C. A. Ellis, A. Sattiraju, R. Miller, and V. Calhoun, “Examining Reproducibility of EEG Schizophrenia Biomarkers Across Explainable Machine Learning Models,” 2022.

[3] W. Samek, G. Montavon, A. Vedaldi, L.K. Hansen, and K.-R. Müller, Eds., Explainable AI: Interpreting, Explaining and Visualizing Deep Learning, vol. 11700. Cham: Springer International Publishing, 2019.

[4] C. A. Ellis et al., “Novel Methods for Elucidating Modality Importance in Multimodal Electrophysiology Classifiers,” bioRxiv, 2022.

[5] S. Pathak, C. Lu, S. B. Nagaraj, M. van Putten, and C. Seifert, “STQS: Interpretable multi-modal Spatial-Temporal-seQuential model for automatic Sleep scoring,” Artif. Intell. Med., vol. 114, no. January, p. 102038, 2021, doi: 10.1016/j.artmed.2021.102038.

[6] C. A. Ellis, R. L. Miller, and V. D. Calhoun, “A Novel Local Explainability Approach for Spectral Insight into Raw EEG-Based Deep Learning Classifiers,” in 21st IEEE International Conference on BioInformatics and BioEngineering, 2021, pp. 0–5.

[7] C. A. Ellis, M. S. E. Sendi, R. Miller, and V. Calhoun, “A Novel Activation Maximization-based Approach for Insight into Electrophysiology Classifiers,” 2021.

[8] C. A. Ellis, R. L. Miller, and V. D. Calhoun, “A Model Visualization-based Approach for Insight into Waveforms and Spectra Learned by CNNs,” in Proceedings of the Annual International Conference of the IEEE Engineering in Medicine and Biology Society, EMBS, 2022, vol. 2022-July, pp. 1643–1646, doi: 10.1109/EMBC48229.2022.9871414.

[9] C. A. Ellis, R. L. Miller, and V. D. Calhoun, “A Systematic Approach for Explaining Time and Frequency Features Extracted by Convolutional Neural Networks From Raw Electroencephalography Data,” Front. Neuroinform., vol. 16, no. May, pp. 1–11, 2022, doi: 10.3389/fninf.2022.872035.

[10] R. Boostani, K. Sadatnezhad, and M. Sabeti, “An efficient classifier to diagnose of schizophrenia based on the EEG signals,” Expert Syst. Appl., vol. 36, no. 3 PART 2, pp. 6492–6499, 2009, doi: 10.1016/j.eswa.2008.07.037.

[11] C. Phang, C. Ting, F. Noman, and H. Ombao, “Classification of EEG-Based Brain Connectivity Networks in Schizophrenia Using a Multi-Domain Connectome Convolutional Neural,” pp. 1–15.

[12] M. A. Vázquez, A. Maghsoudi, and I.P. Mariño, “An Interpretable Machine Learning Method for the Detection of Schizophrenia Using EEG Signals,” Front. Syst. Neurosci., vol. 15, no. May, pp. 1–11, 2021, doi: 10.3389/fnsys.2021.652662.

[13] S.H. Na, S.H. Jin, S.Y. Kim, and B.J. Ham, “EEG in schizophrenic patients: Mutual information analysis,” Clin. Neurophysiol., vol. 113, no. 12, pp. 1954–1960, 2002, doi: 10.1016/S1388-2457(02)00197-9.

[14] W. Mumtaz, L. Xia, M. A. M. Yasin, S. S. A. Ali, and A. S. Malik, “A wavelet-based technique to predict treatment outcome for Major Depressive Disorder,” PLoS One, vol. 12, no. 2, pp. 1–30, 2017, doi: 10.1371/journal.pone.0171409.

[15] R. A. Movahed, G. P. Jahromi, S. Shahyad, and G. H. Meftahi, “A major depressive disorder classification framework based on EEG signals using statistical, spectral, wavelet, functional connectivity, and nonlinear analysis,” J. Neurosci. Methods, vol. 358, no. November 2020, p. 109209, 2021, doi: 10.1016/j.jneumeth.2021.109209.

[16] H. W. Loh, C. P. Ooi, E. Aydemir, T. Tuncer, S. Dogan, and U. R. Acharya, “Decision support system for major depression detection using spectrogram and convolution neural network with EEG signals,” Expert Syst., vol. 39, no. 3, pp. 1–15, 2022, doi: 10.1111/exsy.12773.

[17] S. L. Oh, J. Vicnesh, E. J. Ciaccio, R. Yuvaraj, and U. R. Acharya, “Deep convolutional neural network model for automated diagnosis of Schizophrenia using EEG signals,” Appl. Sci., vol. 9, no. 14, 2019, doi: 10.3390/app9142870.

[18] C. A. Ellis, A. Sattiraju, R. Miller, and V. Calhoun, “Examining Effects of Schizophrenia on EEG with Explainable Deep Learning Models,” 2022.

